## Supplemental Data File 5 for "Rapid response of fly populations to gene dosage across development and generations"

>egfp-bicoid-pLacZattB

gatccactagtgtcgacgatgtaggtcacggtctcgaagccgcggtgcgggtgccagggcgtgcccttgggctccccggg

cgcgtactccacctcacccatctggtccatcatgatgaacgggtcgaggtggcggtagttgatcccggcgaacgcgcggc

gcaccgggaagccctcgccctcgaaaccgctgggcgcggtggtcacggtgagcacgggacgtgcgacggcgtcggcgggt

gcggatacgcggggcagcgtcagcgggttctcgacggtcacggcgggcatgtcgacactagttctagccagcttttgttc

cctttagtgagggttaatttcgagcttggcgtaatcatggtcatagctgtttcctgtgtgaaattgttatccgctcacaa

ttccacacaacatacgagccggaagcataaagtgtaaagcctggggtgcctaatgagtgagctaactcacattaattgcg

ttgcgctcactgcccgctttccagtcgggaaacctgtcgtgccagctgcattaatgaatcggccaacgcgcggggagagg

cggtttgcgtattgggcgctcttccgcttcctcgctcactgactcgctgcgctcggtcgttcggctgcggcgagcggtat

cagctcactcaaaggcggtaatacggttatccacagaatcaggggataacgcaggaaagaacatgtgagcaaaaggccag

caaaaggccaggaaccgtaaaaaggccgcgttgctggcgtttttccataggctccgcccccctgacgagcatcacaaaaa

tcgacgctcaagtcagaggtggcgaaacccgacaggactataaagataccaggcgtttccccctggaagctccctcgtgc

gctctcctgttccgaccctgccgcttaccggatacctgtccgcctttctcccttcgggaagcgtggcgctttctcatagc

tcacgctgtaggtatctcagttcggtgtaggtcgttcgctccaagctgggctgtgtgcacgaaccccccgttcagcccga

ccgctgcgccttatccggtaactatcgtcttgagtccaacccggtaagacacgacttatcgccactggcagcagccactg

gtaacaggattagcagagcgaggtatgtaggcggtgctacagagttcttgaagtggtggcctaactacggctacactaga

aggacagtatttggtatctgcgctctgctgaagccagttaccttcggaaaaagagttggtagctcttgatccggcaaaca

aaccaccgctggtagcggtggtttttttgtttgcaagcagcagattacgcgcagaaaaaaaggatctcaagaagatcctt

tgatcttttctacggggtctgacgctcagtggaacgaaaactcacgttaagggattttggtcatgagattatcaaaaagg

atcttcacctagatccttttaaattaaaaatgaagttttaaatcaatctaaagtatatatgagtaaacttggtctgacag

ttaccaatgcttaatcagtgaggcacctatctcagcgatctgtctatttcgttcatccatagttgcctgactccccgtcg

tgtagataactacgatacgggagggcttaccatctggccccagtgctgcaatgataccgcgagacccacgctcaccggct

ccagatttatcagcaataaaccagccagccggaagggccgagcgcagaagtggtcctgcaactttatccgcctccatcca

gtctattaattgttgccgggaagctagagtaagtagttcgccagttaatagtttgcgcaacgttgttgccattgctacag

gcatcgtggtgtcacgctcgtcgtttggtatggcttcattcagctccggttcccaacgatcaaggcgagttacatgatcc

cccatgttgtgcaaaaaagcggttagctccttcggtcctccgatcgttgtcagaagtaagttggccgcagtgttatcact

catggttatggcagcactgcataattctcttactgtcatgccatccgtaagatgcttttctgtgactggtgagtactcaa

ccaagtcattctgagaatagtgtatgcggcgaccgagttgctcttgcccggcgtcaatacgggataataccgcgccacat

agcagaactttaaaagtgctcatcattggaaaacgttcttcggggcgaaaactctcaaggatcttaccgctgttgagatc

cagttcgatgtaacccactcgtgcacccaactgatcttcagcatcttttactttcaccagcgtttctgggtgagcaaaaa

caggaaggcaaaatgccgcaaaaaagggaataagggcgacacggaaatgttgaatactcatactcttcctttttcaatat

tattgaagcatttatcagggttattgtctcatgagcggatacatatttgaatgtatttagaaaaataaacaaataggggt

tccgcgcacatttccccgaaaagtgccacctgacgcgccctgtagcggcgcattaagcgcggcgggtgtggtggttacgc

gcagcgtgaccgctacacttgccagcgccctagcgcccgctcctttcgctttcttcccttcctttctcgccacgttcgcc

ggctttccccgtcaagctctaaatcgggggctccctttagggttccgatttagtgctttacggcacctcgaccccaaaaa

acttgattagggtgatggttcacgtagtgggccatcgccctgatagacggtttttcgccctttgacgttggagtccacgt

tctttaatagtggactcttgttccaaactggaacaacactcaaccctatctcggtctattcttttgatttataagggatt

ttgccgatttcggcctattggttaaaaaatgagctgatttaacaaaaatttaacgcgaattttaacaaaatattaacgct

tacaatttccattcgccattcaggctgcgcaactgttgggaagggcgatcggtgcgggcctcttcgctattacgccagct

ggcgaaagggggatgtgctgcaaggcgattaagttgggtaacgccagggttttcccagtcacgacgttgtaaaacgacgg

ccagtgaattgtaatacgactcactatagggcgaattgggtacgtaccgggcccctagtatgtatgtaagttaataaaac

ccatttttgcggaaagtagataaaaaaaacattttttttttttactgcactggatatcattgaacttatctgatcagttt

taaatttacttcgatccaagggtatttgatgtaccaggttctttcgattacctctcactcaaaatgacattccactcaaa

gtcagcgctgtttgcctccttctctgtccacagaaatatcgccgtctctttcgccgctgcgtccgctatctctttcgcca

ccgtttgtagcgttacgtagcgtcaatgtccgccttcagttgcattttgtcagcggtttcgtgacgaagctccaagcggt

ttacgccatcaattaaacacaaagtgctgtgccaaaactcctctcgcttcttatttttgtttgttttttgagtgattggg

gtggtgattggttttgggtgggtaagcaggggaaagtgtgaaaaatcccggcaatgggccaagaggatcaggagctatta

attcgcggaggcagcaaacacccatctgccgagcatctgaacaatgtgagtagtacatgtgcatacatcttaagttcact

tgatctataggaactgcgattgcaacatcaaattgtctgcggcgtgagaactgcgacccacaaaaatcccaaaccgcaat

tgcacaaacaaatagtgacacgaaacagattattctggtagctgttctcgctatataagacaatttttgagatcatatca

tgatcaagacatctaaaggcattcattttcgactatattcttttttacaaaaaatataacaaccagatattttaagctga

tcctagatgcacaaaaaataaataaaagtataaacctacttcgtaggatacttcggggtactttttgttcggggttagat

gagcataacgcttgtagttgatatttgagatcccctatcattgcagggtgacagcggagcggcttcgcagagctgcatta

accagggcttcgggcaggccaaaaactacggcacgctccggccacccagtccgccggaggactccggttcagggagcggc

caactagccgagaacctcacctatgcctggcacaatatggacatctttggggcggtcaatcagccgggctccggatggcg

gcagctggtcaaccggacacgcggactattctgcaacgagcgacacataccggcgcccaggaaacatttgctcaagaacg

gtgagtttctattcgcagtcggctgatctgtgtgaaatcttaataaagggtccaattaccaatttgaaactcagtttgcg

gcgtggcctatccgggcgaacttttggccgtgatgggcagttccggtgccggaaagacgaccctgctgaatgcccttgcc

tttcgatcgccgcagggcatccaagtatcgccatccgggatgcgactgctcaatggccaacctgtggacgccaaggagat

gcaggccaggtgcgcctatgtccagcaggatgacctctttatcggctccctaacggccagggaacacctgattttccagg

ccatggtgcggatgccacgacatctgacctatcggcagcgagtggcccgcgtggatcaggtgatccaggagctttcgctc

agcaaatgtcagcacacgatcatcggtgtgcccggcagggtgaaaggtctgtccggcggagaaaggaagcgtctggcatt

cgcctccgaggcactaaccgatccgccgcttctgatctgcgatgagcccacctccggactggactcatttaccgcccaca

gcgtcgtccaggtgctgaagaagctgtcgcagaagggcaagaccgtcatcctgaccattcatcagccgtcttccgagctg

tttgagctctttgacaagatccttctgatggccgagggcagggtagctttcttgggcactcccagcgaagccgtcgactt

cttttcctagtgagttcgatgtgtttattaagggtatctagcattacattacatctcaactcctatccagcgtgggtgcc

cagtgtcctaccaactacaatccggcggacttttacgtacaggtgttggccgttgtgcccggacgggagatcgagtcccg

tgatcggatcgccaagatatgcgacaattttgctattagcaaagtagcccgggatatggagcagttgttggccaccaaaa

atttggagaagccactggagcagccggagaatgggtacacctacaaggccacctggttcatgcagttccgggcggtcctg

tggcgatcctggctgtcggtgctcaaggaaccactcctcgtaaaagtgcgacttattcagacaacggtgagtggttccag

tggaaacaaatgatataacgcttacaattcttggaaacaaattcgctagattttagttagaattgcctgattccacaccc

ttcttagtttttttcaatgagatgtatagtttatagttttgcagaaaataaataaatttcatttaactcgcgaacatgtt

gaagatatgaatattaatgagatgcgagtaacattttaatttgcagatggttgccatcttgattggcctcatctttttgg

gccaacaactcacgcaagtgggcgtgatgaatatcaacggagccatcttcctcttcctgaccaacatgacctttcaaaac

gtctttgccacgataaatgtaagtcttgtttagaatacatttgcatattaataatttactaactttctaatgaatcgatt

cgatttaggtgttcacctcagagctgccagtttttatgagggaggcccgaagtcgactttatcgctgtgacacatacttt

ctgggcaaaacgattgccgaattaccgctttttctcacagtgccactggtcttcacggcgattgcctatccgatgatcgg

actgcgggccggagtgctgcacttcttcaactgcctggcgctggtcactctggtggccaatgtgtcaacgtccttcggat

atctaatatcctgcgccagctcctcgacctcgatggcgctgtctgtgggtccgccggttatcataccattcctgctcttt

ggcggcttcttcttgaactcgggctcggtgccagtatacctcaaatggttgtcgtacctctcatggttccgttacgccaa

cgagggtctgctgattaaccaatgggcggacgtggagccgggcgaaattagctgcacatcgtcgaacaccacgtgcccca

gttcgggcaaggtcatcctggagacgcttaacttctccgccgccgatctgccgctggactacgtgggtctggccattctc

atcgtgagcttccgggtgctcgcatatctggctctaagacttcgggcccgacgcaaggagtagccgacatatatccgaaa

taactgcttgttttttttttttaccattattaccatcgtgtttactgtttattgccccctcaaaaagctaatgtaattat

atttgtgccaataaaaacaagatatgacctatagaatacaagtatttccccttcgaacatccccacaagtagactttgga

tttgtcttctaaccaaaagacttacacacctgcataccttacatcaaaaactcgtttatcgctacataaaacaccgggat

atattttttatatacatacttttcaaatcgcgcgccctcttcataattcacctccaccacaccacgtttcgtagttgctc

tttcgctgtctcccacccgctctccgcaacacattcaccttttgttcgacgaccttggagcgactgtcgttagttccgcg

cgattcggttcgctcaaatggttccgagtggttcatttcgtctcaatagaaattagtaataaatatttgtatgtacaatt

tatttgctccaatatatttgtatatatttccctcacagctatatttattctaatttaatattatgactttttaaggtaat

tttttgtgacctgttcggagtgattagcgttacaatttgaactgaaagtgacatccagtgtttgttccttgtgtagatgc

atctcaaaaaaatggtgggcataatagtgttgtttatatatatcaaaaataacaactataataataagaatacatttaat

ttagaaaatgcttggatttcactggaactaggctagcataacttcgtataatgtatgctatacgaagttatgctagcgga

tccaaaacagaaaatgtcaaattttgtaatttattacccgaatcctgtttctgtcagtgactctttgggtgtttcgcttc

ggcagcggcacacacgggggcattcagcattcagggtcgccgagcggggaagggagaagtcctttgtcacacactgcgtg

tgaatttgtggcccagttccgcacagtggtatctagcagatactttttgtaatatttgtaagaatgcacagaacttcgtt

acattcagagatttctagtttctagtcagatgactaaatacaatattatgtgtcaaagtagttaattttgtttttgaata

aatttctaaaatatcagcttgcctttaacgtaaatatagccttttctacctatctttatctgatataaaaaagtagcatt

tgaaatgctgttgttgcaatggtatcgaattggtattaaagctctggccctgtttaatttgtatttgtagtcagagcaac

cactgtacctttgtgtgtatccttgggctttgcctatgcgaacgtattttgtttgccttttggctgcactccgactccgt

cgactggagtgtctgtgaattgacttttgttgccagttggcagcggcagaagcagcaaagcccggccaacagcaacaagc

tcctgccagatcccaaaagcaaacacgacaattatttggcaaatgtcattaaaaaatatttcacttaaggccttgcgaca

cttgcttaaaggtcaactggctcgttgggtgtgttttaaaatgttaaagcttgggccaatgcactgagcaacttaatgct

tgtagatatttacacaatattcttcaacgctaaacatatcgaattttccaaatatggagcctgaaaataataattgccaa

tcctagcttaaaatcagaaatgagtagaacaacttaaaaaaaattaacaaaagaatcgaacgctacagctaattaactcg

acaactggttaccttttattcttctaatacattttataatgcactgcctaacaggtacatatagcaagcactatatgctg

tcttacaaaacgattatatgatattttctttcgtacgtagccgtttgagatcatttggaaaaacaaactcgatctccacc

atccttattctttgtcccaagtccttatatatctcgcgatactaagattgaataatgtagttattaatagcggaagtatg

taacagaataaactacaaagtgcacattttgttcaattcaggctggactggactggagcatattaatattataatattaa

caaaaattcaaattaaacattcgacacttgtctaattgattcctaaatttggggtgcctgtttgttaattaaatgttaat

attatgaagttccaaacagagcaaagagtttaagtttaattggttctacttatttgttacaatattcaagctttttttta

ttattattctcaaatgcaaatctctacaaataaataaacctccgacgttttagaacattcaccttttgtcagtgagcaca

acctttcaatacagcccgacagggggctctctactgctgtctcttcacgccccctggtgaaaacgctgtgcactcaatcg

gtttgcagctttgccgtactgttcgattaaaaacttttaaattagaggcaaacatttaaaaataaaatgtccaaatattt

gtctaaaatgtattgtagacgcttattgatttttaaattactcaaaagaatgaacatcgagggagggccgccaattgtgc

catctctacatctcttcgctcatccctaaataacggcactctgcagatgcgaagcagtggatcgcaaaaacgcaaaatgt

gggcgaaataagttcgcgagcgtctcgaaagtaaccggttactgaaaatacaagaaagtttccacactcctttgccattt

ttccgcgcggcgcttggaaattcgtaaagataacgcggcggagtgtttggggctagcccaaagatgaaaggagaagaact

tttcactggagttgtcccaattcttgttgaattagatggtgatgttaatgggcacaaattttctgtcagtggagagggtg

aaggtgatgcaacatacggaaaacttacccttaaatttatttgcactactggaaaactacctgttccatggccaacactt

gtcactactttaacttatggtgttcaatgcttttcaagatacccagatcatatgaaacagcatgactttttcaagagtgc

catgcccgaaggttatgtccaggaaagaactatatttttcaaagatgacgggaactacaagacacgtgctgaagtcaagt

ttgaaggtgatacccttgttaatagaatcgagttaaaaggtattgattttaaagaagatggaaacattcttggacacaaa

ttggaatacaactataactcacacaatgtatacatcatggcagacaaacaaaagaatggaatcaaagttaacttcaaaat

tagacacaacattgaagatggaagcgttcaactagcagaccattatcaacaaaatactccaattggcgatggccctgtcc

ttttaccagacaaccattacctgtccacacaatctgccctttcgaaagatcccaacgaaaagagagaccacatggtcctt

cttgagtttgtaacagctgctgggattacacatggcatggatgaactatacaaagcatgcggtggttctggtggtggttc

tggtatggcgcaaccgccgccagatcaaaacttttaccatcatccgctgccccacacgcacacacatccgcatccgcact

cccatccgcatccgcactcgcatccgcacccacatcaccaacatccgcagcttcagttgccgccacaattccgaaatccc

ttcgatttggtgagttcccatcgcagcagagaagggctcttgtcccaggaaagctacagtacagattccctatggtgaac

aaacaaccagtgcgatcactgatgaccataaacatttattgagccgcagcaaatgtgtttctagaacatagggcgaaatc

ttctattatcttgtttgtgacttttaaagtatcgtagcagaatctaaaaaccaattgatattattaatcgttacagttag

tatagtatataattgtatatgaattgtggggcatcatgttattagtgatttgccgaaatgttctaaaaggtgtttcattg

aaatggacgaatgttaaacctgttgcactcacaccgacaatcagtaatgtctatttttcaaaagccacatctatggccac

tgggtatacattattgactttatacacttcatacaacatattttctaaaacaagcattgttgtcctgcatgatgattagt

gaaagtaatattgcaagattcggtccccgaagcgaatcgtcctttcacgtttttatataaagacagtgtaccccttgatt

ctttgaagcttttcgatgagcgaacgggagcgataaactacaactacatacgtccgtatctgcccaaccagatgcccaag

ccaggtgagctcaaagccaacaaagtcagccatcgtcttatcagatgtctttccctcagaggagctgcccgactctctgg

tgatgcggcgaccacgtcgcacccgcaccacttttaccagctctcaaatagcagagctggagcagcactttctgcaggga

cgatacctcacagccccccgacttgcggatctgtcagcgaaactagccctgggcacagcccaggtgaagatatggtttaa

gaaccgtcggcgtcgtcacaagatccaatcggatcagcacaaggaccagtcctacgaggggatgcctctctcgccgggta

tgaaacagagcgatggcgatccccccagcttgcagactcttagcttgggtggaggagccacgcccaacgctttgactccg

tcacccacgccctcaacgcccactgcacacatgacggagcactacagcgagtcattcaacgcctactacaactacaatgg

aggccacaatcacgcccaggccaatcgtcacatgcacatgcagtatccttccggaggggggccaggacctgggtcgacca

atgtcaatggcggccagttcttccagcagcagcaggtccataatcaccagcagcaactgcaccaccagggcaaccacgtg

ccgcaccagatgcagcagcagcaacagcaggctcagcagcagcaataccatcactttgacttccagcaaaagcaagccag

cgcctgtcgcgtcctggtcaaggacgaaccggaggccgactacaacttcaacagctcgtactacatgcgatcgggaatgt

ctggcgccactgcatcggcatccgctgtggcccgaggcgctgcctcgccgggctccgaggtctacgagccattaacaccc

aagaatgacgaaagtccgagtctgtgtggcatcggcatcggcggaccttgcgccatcgccgttggcgagacggaggcggc

cgacgacatggacgacggaacgagcaagaagacgacgctacaggtcaggcatgagtccacaaccttttttgatctcttga

ttctgagtgtggcgtttataaattgaagctttaagctttgtaactttcaaactgtctggtttgagatgttattctgaaag

tacttctatttccgatcgatgagatttgggagttctccaatatttaacatttaacttattaagtttttgttttctaaatt

agacatggcatttctgaaagggaagtacaagtgttaaagatgtattttaatatagaatttgtatcaaaggttaagatttc

aaccgtttgaaagcccttagttttcagggttttttacttttttattcatgtaatcactcttaatacactgcaagttaaaa

tagcatttctttgaccagaaaaataagatctatgcattttaaaagtgaaaacagactcatatgctgatgaacatttttag

ctataaattgtaacaataatttagcaatttcaatcgaatttatttatgttctaaatgcgttcgctctctccctagatctt

ggagcctttgaagggtctggacaagagctgcgacgatggcagtagcgacgacatgagcaccggaataagagccttagcag

gaaccggaaatcgtggagcggcatttgccaaatttggcaagccttcgcccccacaaggccctcagccgcccctcggaatg

gggggcgtggccatgggcgaatcgaaccaatatcaatgcacgatggatacgataatgcaagcgtataatccccatcggaa

cgccgcgggcaactcgcagtttgcctactgcttcaattagcctggacgagaggcgtgttagagagtttcattagctttag

gttaaccactgttgttcctgattgtacaaataccaagtgattgtagatatctacgcgtagaaagttaggtctagtcctaa

gatccgtgtaaatggttcccagggaagttttatgtactagcctagtcagcaggccgcacggattccagtgcatatcttag

tgatactccagttaactctatactttccctgcaatacgctattcgccttagatgtatctgggtggctgctccactaaagc

ccgggaatatgcaaccagttacatttgaggccatttgggcttaagcgtattccatggaaagttatcgtcccacatttcgg

aaattatattccgagccagcaagaaaatcttctctgttacaatttgacatagctaaaaactgtactaatcaaaatgaaaa

atgtttctcttgggcgtaatctcatacaatgattacccttaaagatcgaacatttaaacaataatatttgatatgatatt

ttcaatttctatgctatgccaaagtgtctgacataatcaaacatttgcgcattctttgaccaagaatagtcagcaaattg

tattttcaatcaatgcagaccatttgtttcagattctgagattttttgctgccaaacggaataactatcatagctcacat

tctatttacatcactaagaagagcattgcaatctgttaggcctcaagtttaattttaaaatgctgcacctttgatgttgt

ctctttaagctttgtatttttaattacgaaaatatataagaactactctactcgggtaaattgtgactaactacacataa

ctacatacttagcccatatttccgtccctttctagaatgaacgaaaacagtatctggttttcccgaaaatcttatgaatt

taaaaatgcactttattgcacatactcacacatgcctgccataaaatatgattcgcgatttttccgcgaacacccgcgga

tcataaaacatttgcaccagctgcctgtgtttattcacctacctgaaacccatactcttatcgcctgatcctcgcgcggt

cgcactatttaggtagacactgtacaggcagcactagcggggcgccgccaggcaacttgttgcccctattttctttgata

taagttttgggaaatccagaagtccaataacggggcctatatgaacgaatattgattgggtttctcccagttatagagtt

tttattgatcttgggcgcgttttagtaggtggattatggatgtcctgggaaaaccaagctctgaatccattcttacttat

cacaagacttgttttcctgtgcccttcatcacacctagtttaactactataattaataaactactttttaatccgccatt

tatgaagataatttatagcatctatgcatctttcaaatatcgcgagatatcgcaaattattcataaattagttttttatt

gatattgtaaagctttttaatttggttcaggtttttattgtaaatacttttcacaattgtgcacagtgacccattaacat

ggttgcgatattaacttaagacttaaacaagttaattaagagttttagcacttcttttcaaaagatcatgttttaattat

gtttttaaatgaatcttttcgacaattctaaaaatattatttcgaaaatgttaattttcatatataacctaagtaaagat

attttcaactgtacctcactctcgccgcgacatttccacacggttatcctgttatgacttttcctttgttccaacgtatt

tttagcg

egfp, bicoid upstream and downstream sequence, bicoid gene, NheI, SphI
