## Supplemental figures and tables for "Rapid response of fly populations to gene dosage across development and generations"

Supplementary Fig. 1.

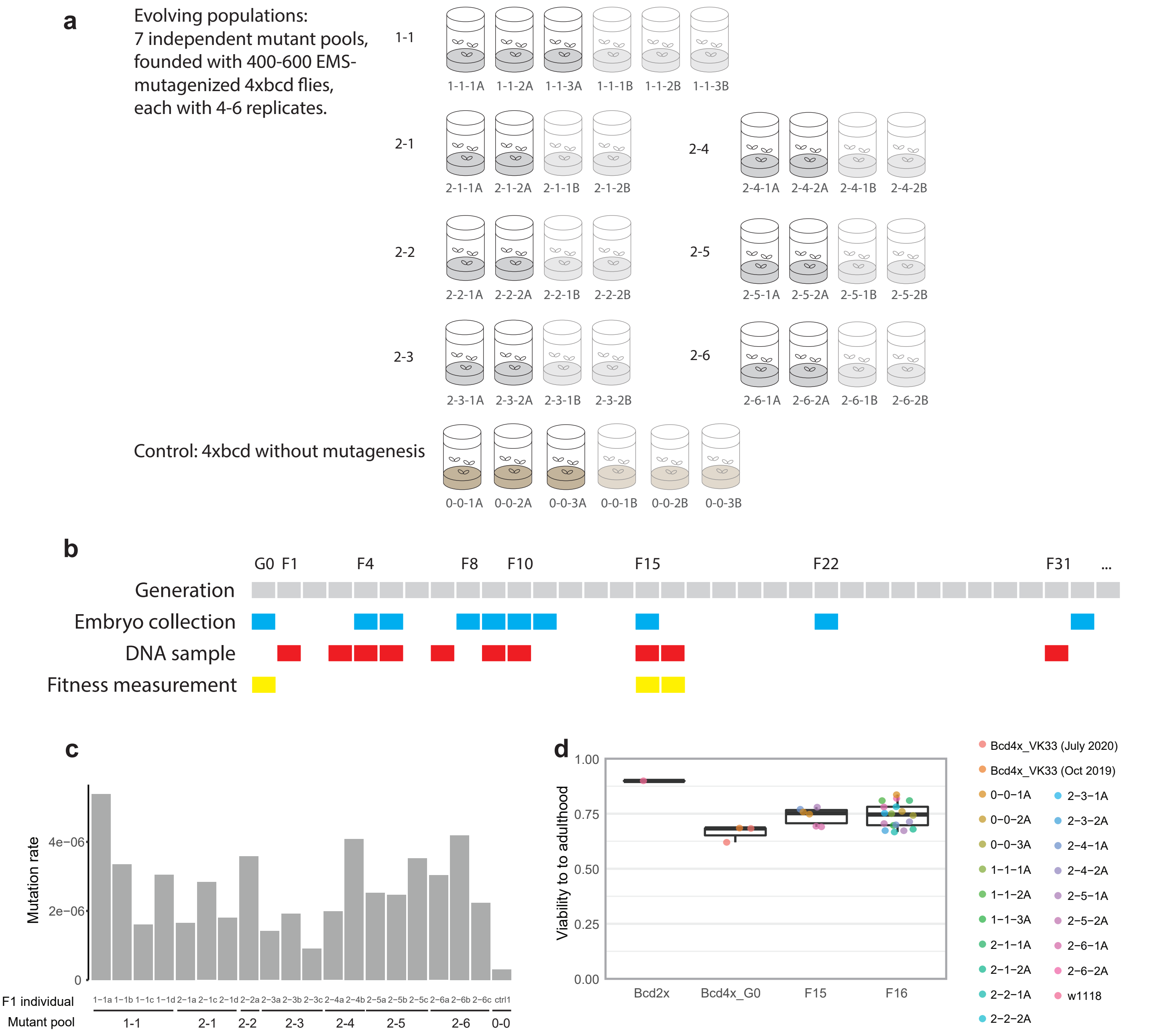


**Supplementary Fig. 1. Mutagenesis, experimental evolution and sampling scheme.** (**a**) The evolving populations were from 7 independent mutant pools each with 4-6 replicates. 4x*bcd* flies without mutagenesis were evolved in parallel as control for standing variation in the lab stock. (**b**) Sampling time for embryo collection, DNA and fitness measurement. (**c**) Mutation rate of 19 mutagenized F1 individuals from the seven mutant pools and one individual from the background population, measured by the number of mutations divided by number of bases covered by whole-genome sequencing in each invidual. (**d**) Viability to adulthood of the evolving populations, compared to wild-type (Bcd2x, w1118) and the ancestral stock (Bcd4x_VK33, measured at two time points).

Supplementary Fig. 2.

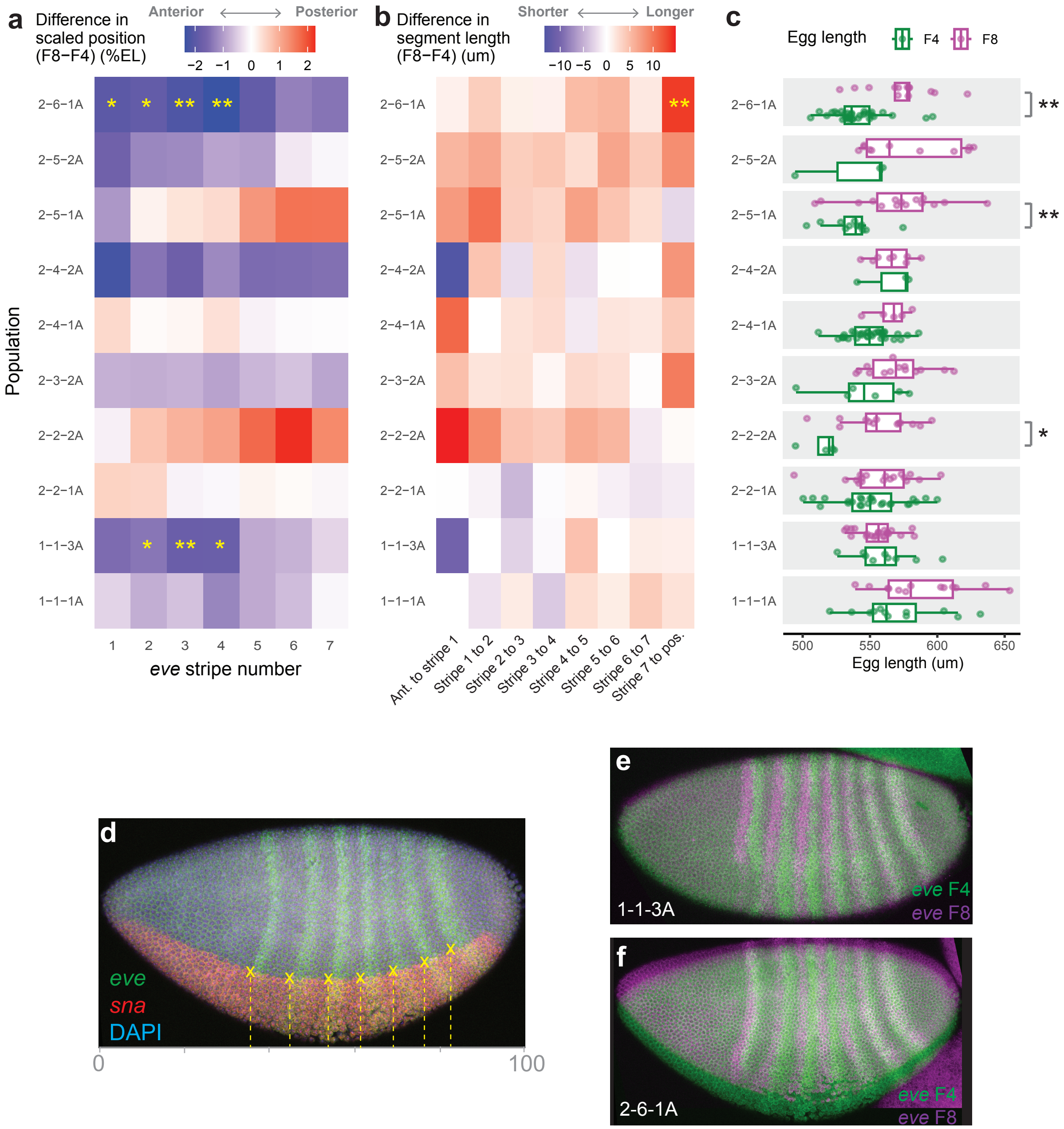


**Supplementary Fig. 2. Response of different populations.** (**a**) Heatmap shows the difference in average *eve* stripe positions between F4 and F8 in each population. (**b**) The difference in average inter-stripe distance between generations. Asterisks in (**a**) and (**b**) indicate statistically different distributions between F4 and F8 based on Wilconxon tests with FDR-correction. (**c**) Distribution of egg length of F4 and F8 embryos, including embryos from (**a**) and (**b**) as well as additionally imaged embryos. Asterisks indicate p-values from Wilcoxon tests. *, p<0.05; **, p < 0.01. (**d**) A representative embryo, showing that the *eve* stripe positions were measured at the intersection between *eve* and *sna* expression, detected by *in situ* hybridization. (**e-f**) Representative embryos from 1-1-3A (**e**) and 2-6-1A (**f**), with F4 and F8 embryos scaled together.

Supplementary Fig. 3.

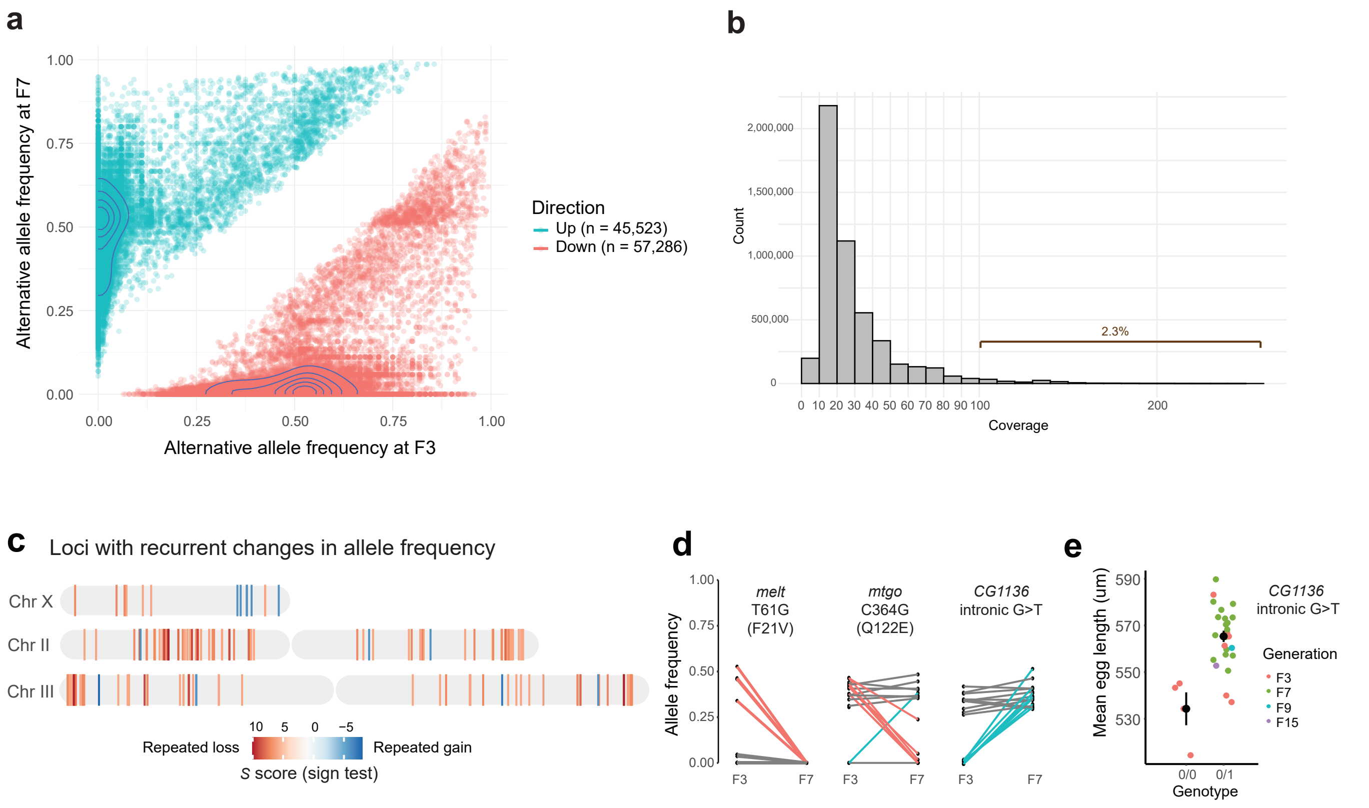


**Supplementary Fig. 3. Changes in allele frequency in evolved populations.** (**a**) Alternative allele frequency at Generation 3 (F3) and Generation 7 (F7). Each point represents a comparison between two generations for a given variant in a given population. 102,809 significant changes on 54,045 sites across 18 populations (Fisher's exact test, FDR-adjusted $p < 0.05$) are shown. Colors represent the direction of changes (blue, increase in alternative allele frequency; red, decrease in alternative allele frequency). Contour lines mark high-density areas on the plot, showing that most changes were transitions between heterozygous (allele frequency around 0.5) and homozygous (allele frequency of 1) states. (**b**) Distribution of sequencing depth on all the variants surveyed with Fisher's test (450,739 variants across two generations and 18 populations). The median was 21 and the mean was 29. (**c**) Variants with consistent increased (blue) or decreased (red) allele frequency in six or more populations, with examples in (**d**). Colored lines indicate significant changes based on Fisher's exact test (red, decrease; blue, increase, FDR-adjusted $p<0.05$). (**e**) Genotype of the SNP in *CG1136* was correlated with embryo length (p=8.36e-05). Black points represent the average phenotype of populations with the same genotype, and error bars represent standard errors.

Supplementary Fig. 4.

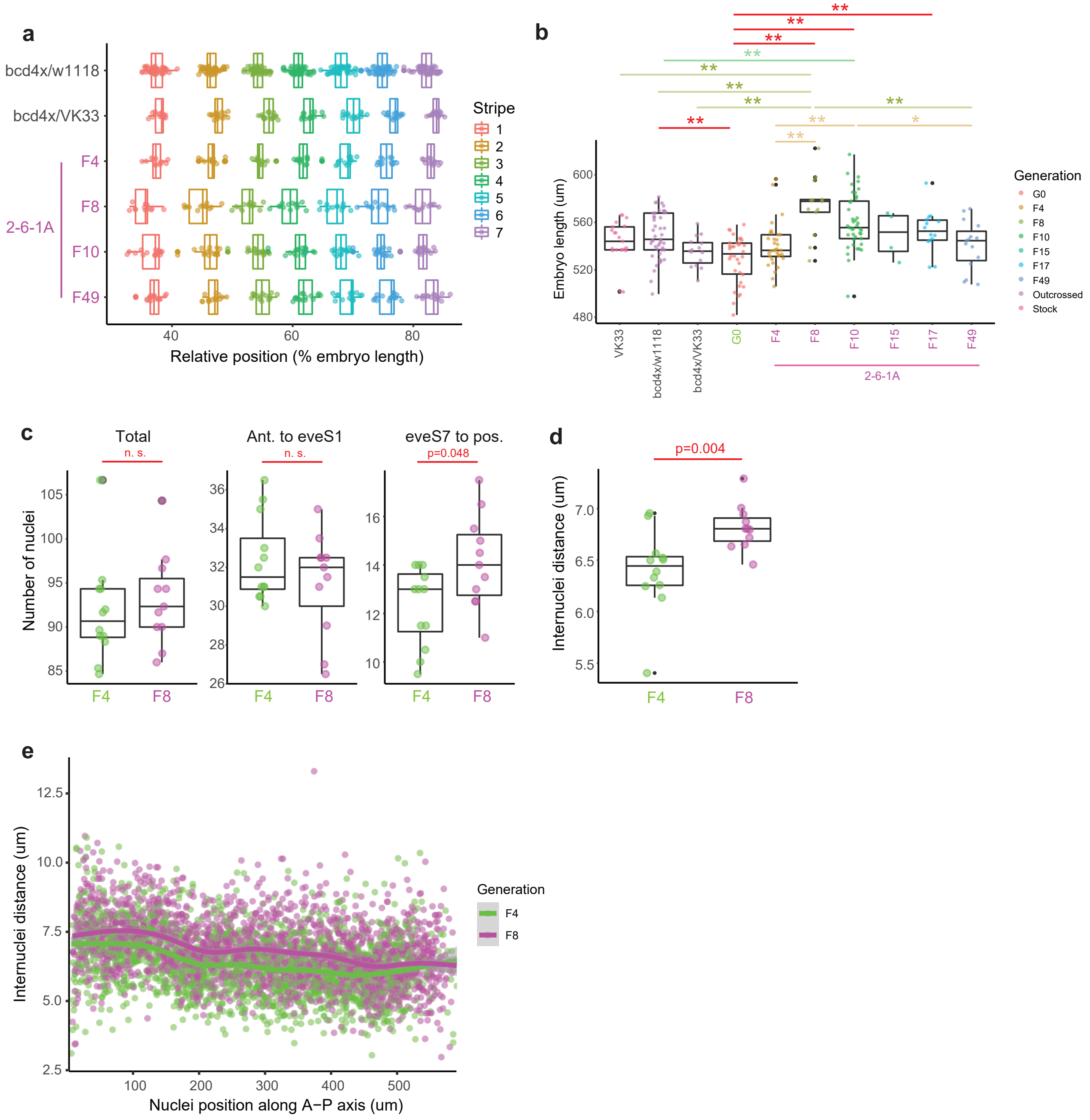


**Supplementary Fig. 4. Embryonic phenotypes of the evolved line 2-6-1A.** *eve* stripe positions (**a**) and embryo length (**b**) changed across time. “bcd4x/w1118” and “bcd4x/VK33” are 4xbcd lines established from outcrosses between 2-6-1A (Generation 40) and wild-type w1118 or VK33, to represent a baseline phenotype without any evolutionary changes. Asterisks in (**b**) represent p-values from pairwise comparisons using Tukey’s test. *, p<0.05. **, p < 0.01. (**c**) Number of nuclei along the *sna* border across the A-P axis. For each embryo, counts from two independent experimenters were averaged (the counts from the two experimenters were not significantly different). Particularly, the data of total number of nuclei were averaged across two measurements by X.C.L. and one measurement by L.G. Other data were only measured once by each experimenter. (**d**) Average inter-nuclei distance per embryo. The data were averaged across all nuclei counted along the A-P axis and across two experimenters’ measurements for each embryo. P-values in (**c**) and (**d**) are from Wilcoxon tests. n. s., not significant. (**e**) Inter-nuclei distance (the 2D-distance between the center of a given nuclei and that of its neighboring nuclei at the anterior side) as a function of nuclei position along the A-P axis, with data from both experimenters. Trend lines were generated with generalized additive mode smoothing (“gam” method), showing that F8 embryos had larger nuclei in the anterior and in the middle.

Supplementary Fig. 5.

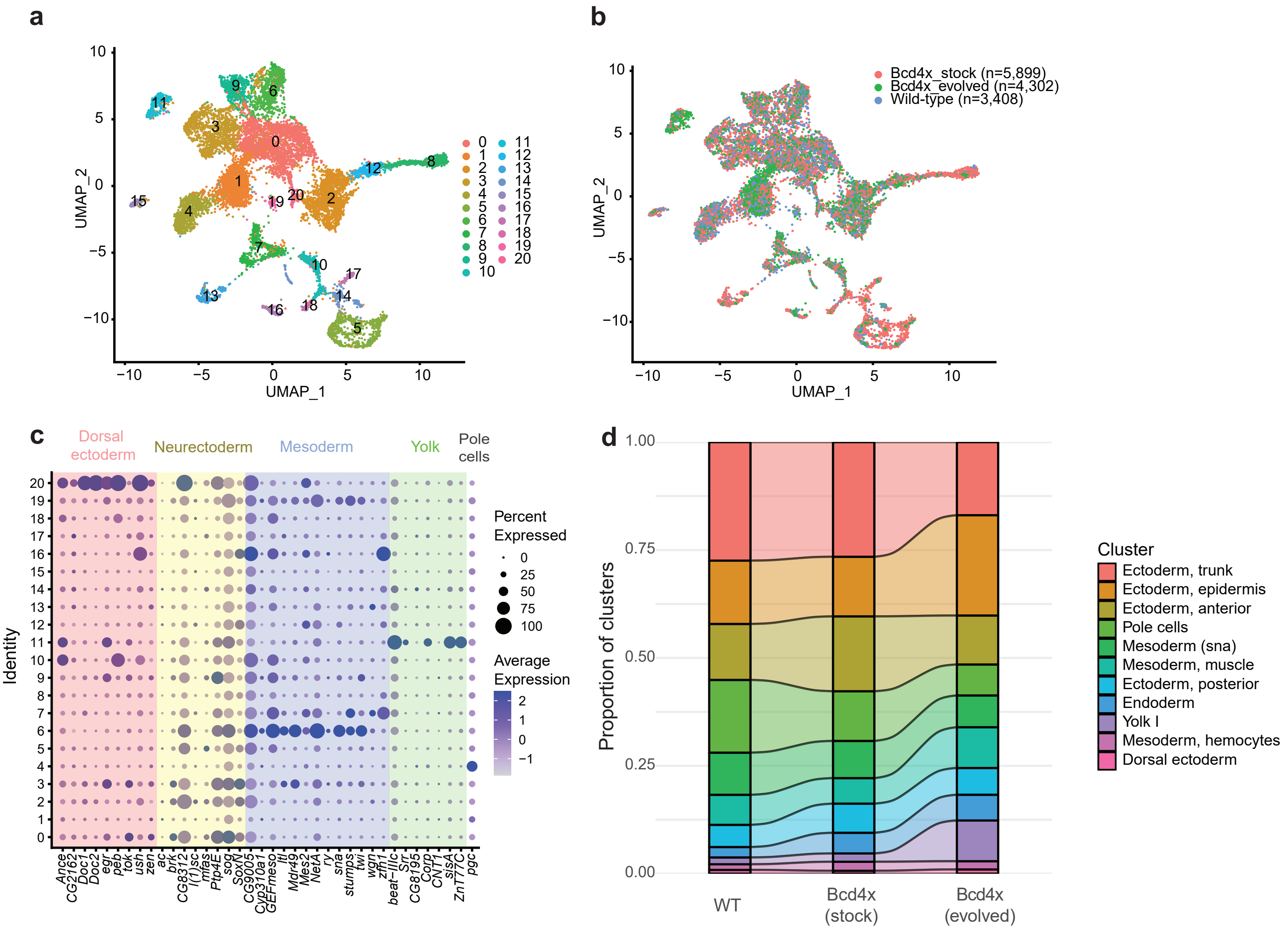


**Supplementary Fig. 5. Single-nuclei transcriptomes of the evolved line 2-6-1A (Generation 20). (a-b)** UMAP of single-nuclei transcriptomes across three samples, with colors representing cell types in (**a**) and sample identity in (**b**). n represents the number of nuclei in each sample. Wild-type is VK33. (**c**) Dot plot showing expression of marker genes for primordium assignment used in Karaiskos, Wahle et al. (2017). (**d**) Proportion of clusters in three samples, only including early embryonic cell types (see **Extended Data Table 2**).

Supplementary Fig. 6.

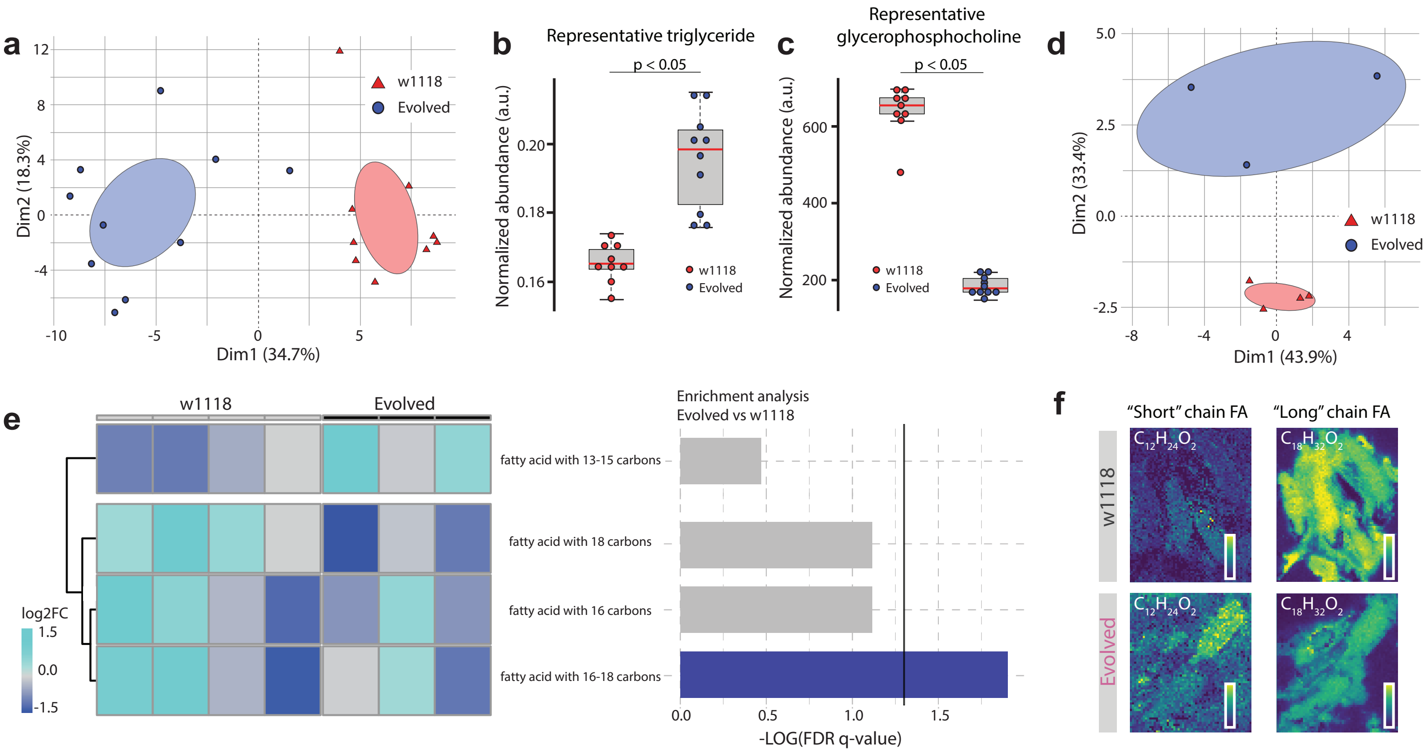


**Supplementary Fig. 6. Metabolic alterations in oocytes from the evolved line 2-6-1A.**
(**a**) Principal Components Analysis (PCA) based on single-oocyte abundance values for 122 different lipids identified by MALDI imaging mass spec. Each dot represents an individual oocyte. n=9 for w1118, n=10 for 2-6-1A. The ellipses delimit the 95% confidence interval.
(**b** & **c**) Abundance values of a representative triglyceride (**b**) and glycerophosphocholine (**c**) from single oocytes obtained through MALDI-imaging mass spec. Each dot represents an individual oocyte with n=9 for w1118 and n=10 for 2-6-1A. The boxed region is one s.d. and the tails are two s.d. (95 %). $p<0.05$, Student's T-test.
(**d**) Principal Components Analysis (PCA) based on single-oocyte abundance values for 21 different fatty acids identified through tandem mass spectrometry followed by MALDI imaging. Each dot represents an individual oocyte. n=4 for w1118, n=3 for 2-6-1A. The ellipses delimit the 95% confidence interval.
(**e**) Relative abundance of fatty acids (left panel, rows) across oocytes (left panel, columns) and their enrichment (right panel), based on 21 fatty acids detected through tandem mass spectrometry coupled with MALDI-imaging. The vertical line represents a threshold of q=0.05.
(**f**) Spatial distribution of a representative C12 (m/z= 199.17031; left) or C18 fatty acid (m/z= 269.24856; right) in sectioned ovaries from the w1118 and 2-6-1A lines.

Supplementary Fig. 7.

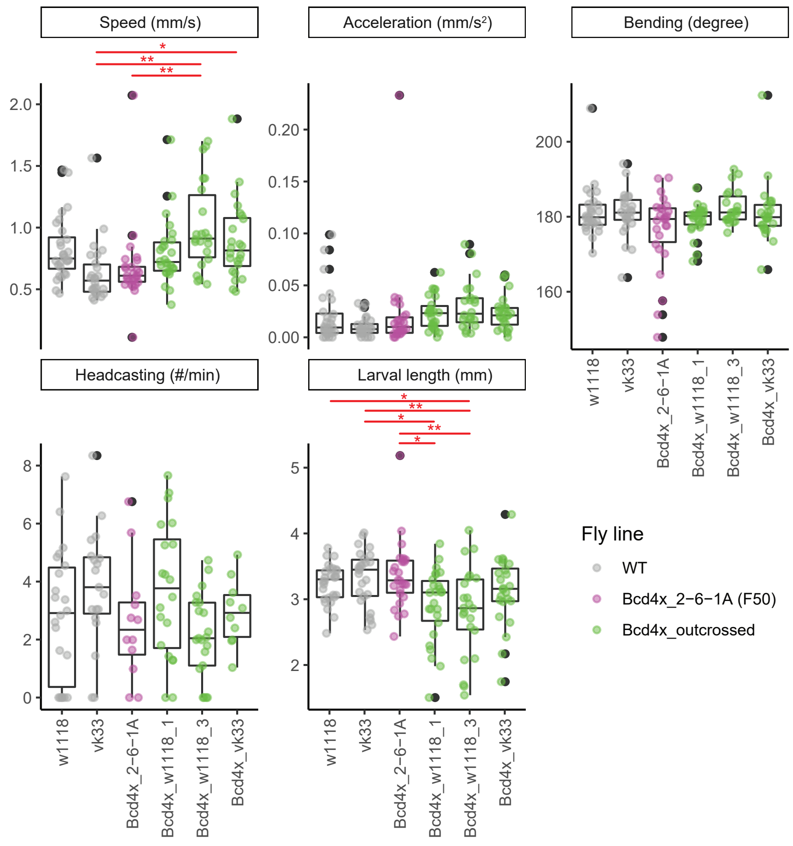


**Supplementary Fig. 7. Quantification of crawling behavior of 3rd-instar larvae from 2-min videos.** Laval length was also extracted from the videos. The evolved line 2-6-1A was from Generation 50. Bcd4x_w1118_1 and Bcd4x_w1118_3 are two biological replicates of Bcd4x lines, outcrossed to the w1118 background. Bcd4x_vk33 is the bcd4x line outcrossed to VK33 background (see **Methods** and **Extended Data Fig. 4** for details of the outcrosses). Asterisks indicate p-values from pair-wise comparisons using Tukey's test. *, $p < 0.05$. **, $p<0.01$. The evolved line did not show any consistent differences from the wild-type or the outcrossed lines.

Supplementary Fig. 8.

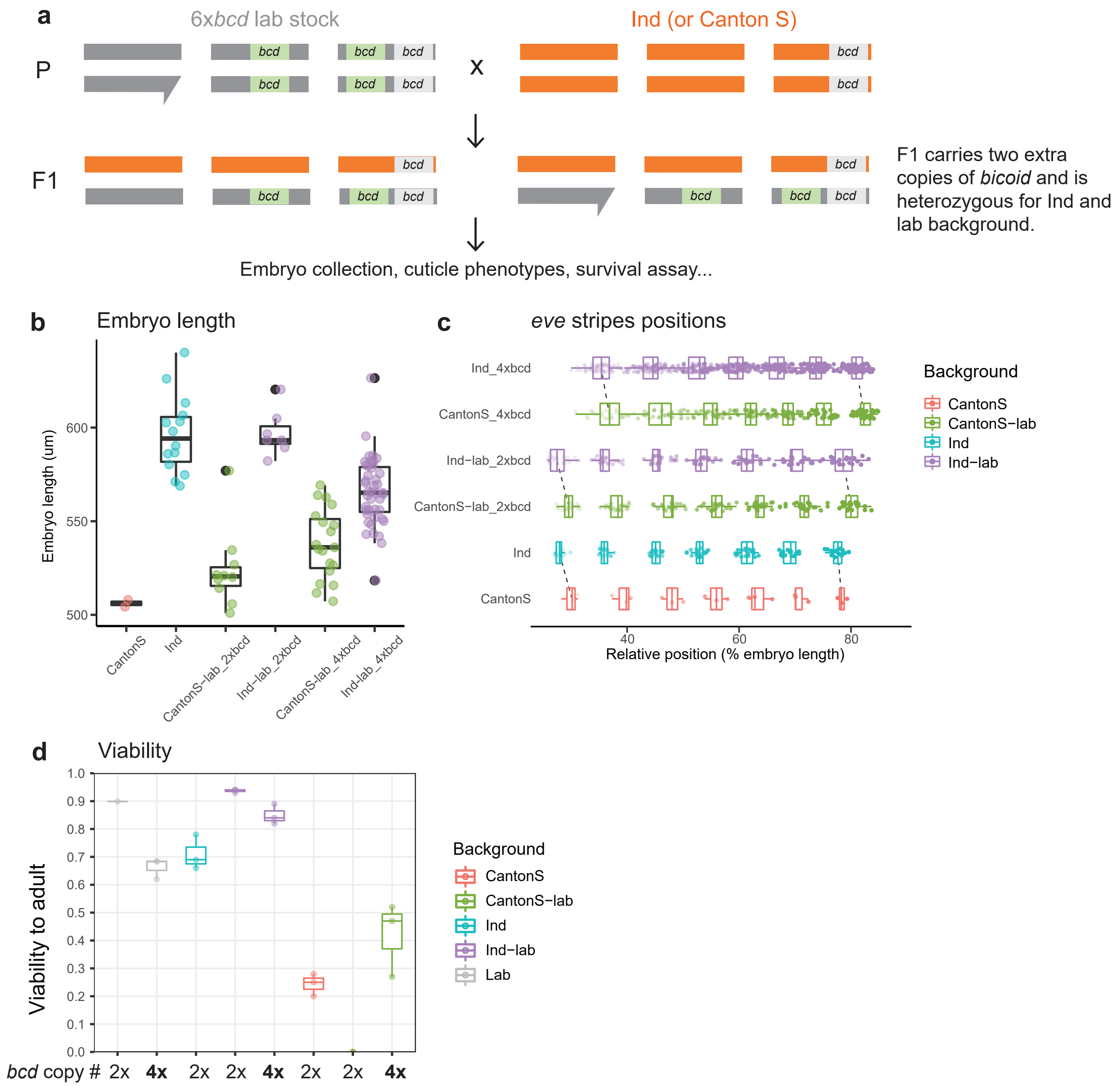


**Supplementary Fig. 8. Cross bicoid transgenes into Ind and Canton-S.** (**a**) The crossing scheme. The dark grey and orange shapes represent the X/Y, 2nd and 3rd chromosomes in 6x*bcd* lab stock and wild strains, respectively. The endogenous copy of *bicoid* and the *egfp-bicoid* transgenes are shown in grey and green boxes, respectively. The position of *bicoid* copies was not to scale. (**b**) Embryo length of Canton-S, Ind, and the F1 flies from crossing the two lines to a lab strain VK33 (CantonS-lab_2xbcd, Ind-lab_2xbcd) or a 6x*bcd* line (CantonS-lab_4xbcd, Ind-lab_4xbcd). (**c**) *eve* stripe positions, detected by *in situ* hybridization. The *eve* stripes of Ind-derived lines were located further to the anterior than those of Canton-S-derived lines under both *bicoid* dosages.  (**d**) Viability to adult of the above-mentioned lines, along with the lab strains (from the data in **Table S1**, with the 4x*bcd* line in VK33 background). CantonS-lab (2x*bcd*) embryos had an extremely low viability, probably due to genomic incompatibility between VK33 and Canton-S.

Supplementary Fig. 9.

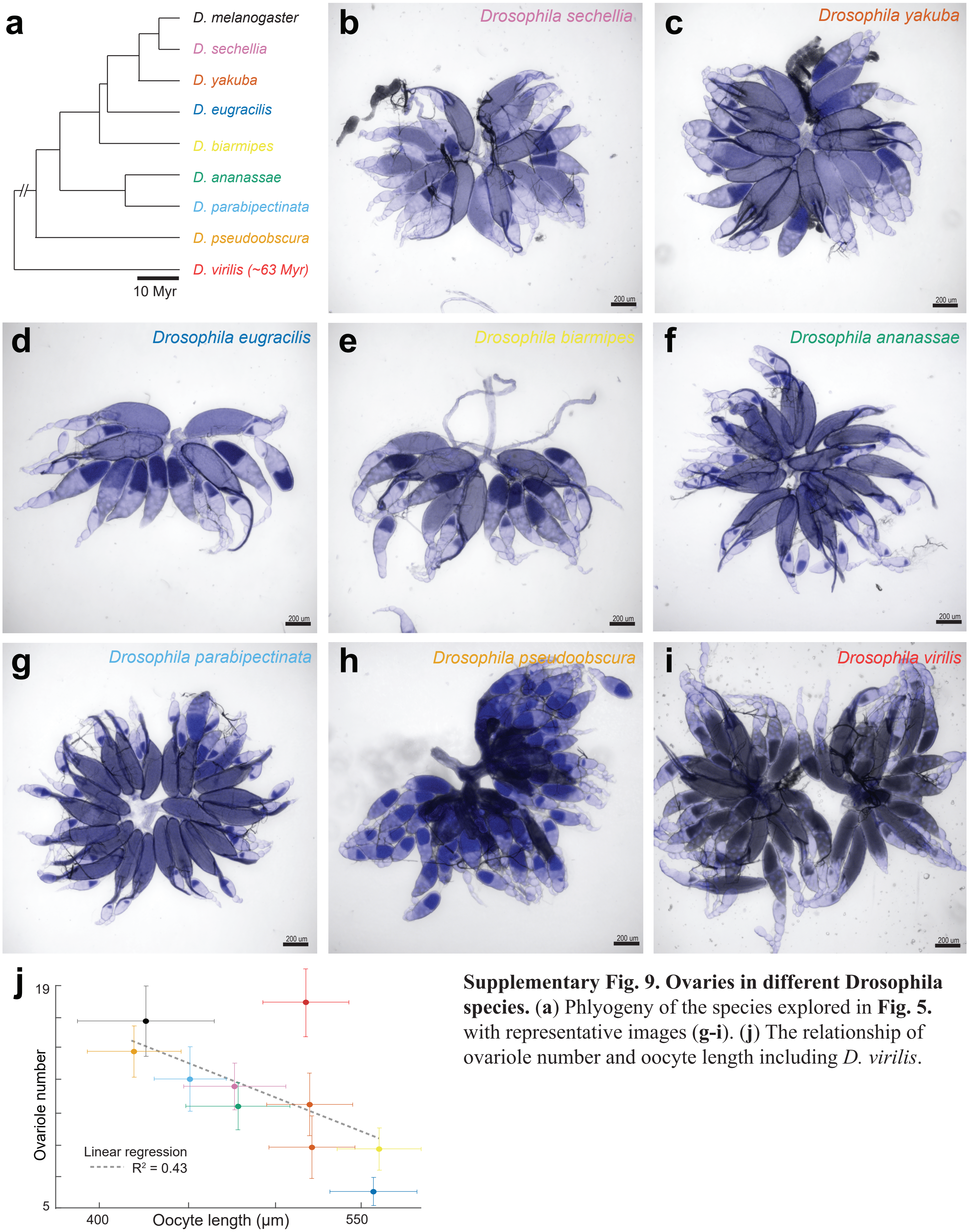


**Supplementary Fig. 9. Ovaries in different Drosophila species.** (**a**) Phlyogeny of the species explored in **Fig. 5.** with representative images (**g-i**). (**j**) The relationship of ovariole number and oocyte length including *D. virilis*.

**Table S1. Viability of stocks carrying 2x-to-6x-*bicoid* (prior to selection).**

| Stock | Dosage of *bicoid* | Hatch rate (24h) | Viability to adult |
|---|---|---|---|
| **w1118** | 2x | 0.73 | 0.899 |
| **4xbcd_VK18** | 4x | 0.68 | 0.853 |
| **4xbcd_VK33** | 4x | 0.72 | 0.685 |
| **6xbcd** | 6x | 0.49 | 0.389 |

**Table S2. Marker genes and cell types for clusters in single nuclei RNA-seq.**

| Cluster | Selected top marker genes | Inferred cell type | Early embryonic cells (stage 4-6)? | % of total nuclei (across three samples) |
|---|---|---|---|---|
| 0 | *lov, opa, hth* | Ectoderm, trunk | Y | 15.6 |
| 1 | *Ribosomal genes* | Cytosolic debris | | 14.4 |
| 2 | *ab, ds, ovo, heph, aop, ft, Wdr62* | Ectoderm, related to epidermis | Y | 10.9 |
| 3 | *oc, knrl, Lim1, jigr1* | Ectoderm, anterior | Y | 9.3 |
| 4 | *pgc, exu, gnu, nos* | Pole cells | Y | 7.6 |
| 5 | *nahoda, verm, exp* | Tracheal system | | 6.0 |
| 6 | *sna, GEFmeso, twi* | Mesoderm | Y | 5.6 |
| 7 | *CadN, zfh1, Him, meso18E, tin* | Mesoderm, related to muscle | Y | 4.7 |
| 8 | *Rdl, para, nAChRalpha6, dysc, slo* | Neuron | | 4.6 |
| 9 | *byn, fkh, tll, disco* | Ectoderm, posterior | Y | 4.0 |
| 10 | *peb, GATAe* | Endoderm | Y | 2.9 |
| 11 | *ZnT77C, sisA, beat-IIIc* | Yolk I | Y | 2.6 |
| 12 | *scrt, wor, insc, nerfin-1, ase, insb, stan, elav* | Central nervous system | | 2.1 |
| 13 | *bt, Mhc, sls* | Muscle | | 2.1 |
| 14 | *AhcyL2, su(r), Gs2, CG9932, Apoltp, Tps1* | Yolk II | | 2.0 |
| 15 | *bcd, CG14229, CG6418, TfIIFbeta, GCC185, GFP* | Unknown | | 1.2 |
| 16 | *Pxn, CG4829, Sr-Cl, ham, vri, gcm2, ct, pnr, RhoL, Pvr, zfh1, gcm, C1GalTA* | Mesoderm, precursors of hemocytes and lymph glands | Y | 1.2 |

| | | | | |
|---|---|---|---|---|
| 17 | *prc, Apoltp, Hayan, Tig, CG10026, fon, daw, vkg, CG9932, AkhR, CG7149, svp, CG2736* | Fat body | | 1.1 |
| 18 | *CG4839, CG11453, CG8907, CG8925, Vha100-5, CG4301, Drip, wdp, Traf-like* | Intestine and Malpighian tubules | | 0.9 |
| 19 | *svp, CG15646, CG10663, HGTX, CG1673, CG7191, htl, ham* | Mesoderm, related to glia development | | 0.8 |
| 20 | *C15, pnr, Doc2, peb, Doc3, tup, ush, Doc1* | Dorsal ectoderm, amnioserosa | Y | 0.5 |
